# MOPower: an R-shiny application for the simulation and power calculation of multi-omics studies

**DOI:** 10.1101/2021.12.19.473339

**Authors:** Hamzah Syed, Georg W Otto, Daniel Kelberman, Chiara Bacchelli, Philip L Beales

## Abstract

**Background:** Multi-omics studies are increasingly used to help understand the underlying mechanisms of clinical phenotypes, integrating information from the genome, transcriptome, epigenome, metabolome, proteome and microbiome. This integration of data is of particular use in rare disease studies where the sample sizes are often relatively small. Methods development for multi-omics studies is in its early stages due to the complexity of the different individual data types. There is a need for software to perform data simulation and power calculation for multi-omics studies to test these different methodologies and help calculate sample size before the initiation of a study. This software, in turn, will optimise the success of a study.

**Results:** The interactive R shiny application MOPower described below simulates data based on three different omics using statistical distributions. It calculates the power to detect an association with the phenotype through analysis of n number of replicates using a variety of the latest multi-omics analysis models and packages. The simulation study confirms the efficiency of the software when handling thousands of simulations over ten different sample sizes. The average time elapsed for a power calculation run between integration models was approximately 500 seconds. Additionally, for the given study design model, power varied with the increase in the number of features affecting each method differently. For example, using MOFA had an increase in power to detect an association when the study sample size equally matched the number of features.

**Conclusions:** MOPower addresses the need for flexible and user-friendly software that undertakes power calculations for multi-omics studies. MOPower offers users a wide variety of integration methods to test and full customisation of omics features to cover a range of study designs.

## Background

In recent years, multi-omics studies, i.e. the integration of data sets from different classes of functionally relevant molecules (e.g. DNA, RNA and protein), have emerged as a valuable approach to provide in depth molecular phenotype information, primarily due to the advancements of high throughput technologies. Large genetic sequencing programs for Mendelian diseases are underway internationally, but these only address the genomic landscape in a single dimension; that of defining disease-causing variants. Other single omics approaches can provide lists of differentially expressed transcripts or proteins that may be associated with the particular condition under investigation, which can be useful in identifying biomarkers and understanding the biological pathways involved in disease pathogenesis. However, the vast majority of genes and their products function as part of complex biological networks with other gene products. Analysis of only one data type is limited to correlations, mostly reflecting reactive processes rather than causative ones which is insufficient for a full understanding of gene function [1]. Integration of multi-omic information from multiple sources including the genome, transcriptome, metabolome, epigenome, microbiome and proteome achieves a broader view of the flow of information for a more reliable understanding of biological processes. This facilitates a better understanding of the molecular changes associated with disease pathogenesis, with the ultimate goal to identify therapeutic targets or clinically relevant biomarkers. For example, Li et al. [2] demonstrate the improvement in accuracy of molecular classification of COPD in small cohorts using mRNA, miRNA, proteomics and metabolomics data from available biological samples. In practice, however, availability of molecular data is often limited by the restricted access to biological samples, which in the case of rare diseases have to be ethically sourced from patients – typically excess samples that are collected for clinical or diagnostic purposes. A careful design of such an investigation, including a rigorous power analysis, is key to successfully deal with the limited resources available.

Analysing high-dimensional and heterogeneous multi-omics data sets poses a number of challenges: (i) Identification of methods that work best for each given single omics dataset, (ii) identification of methods that can integrate, identify and decipher the relevant information from a combination of different data sets and (iii) optimisation of a study’s success by calculating power and sample size. Even with the relatively recent inception of multi-omics studies, several integration methods have been developed each with their own benefits, caveats and the class of information they provide (extensively reviewed by Huang *et al*. [3]). Popular methods can be classified as regression-based, machine learning, Bayesian, matrix factorisation or network analysis. Power and sample size calculation is critical to maximising the success of a study as underpowered studies will produce false-positive associations, miss true signals and can be costly without any benefit. The power to detect associations depends on a multitude of factors that are difficult to account for, especially when each omics dataset has unique characteristics. Therefore the power of analytical models also needs to be tested to identify which model is suitable for the data. Power calculators exist at the individual omics levels for many different outcomes. In genomics, GAS [4] and survivalGWAS_Power [5] are widely used software tools. For RNA-Seq data, there is the R package PROPER [6] and the web tool RNAseqPS [7]. Outside of packaged software, several reports discuss the framework for simulating and calculating sample size and power for RNA-Seq experiments [8–11]. None of these tools can be applied to multi-omics studies, therefore to address this need we have developed the R shiny application MOPower. MOPower simulates multi-omic data (genome, transcriptome and epigenome) and offers a choice of analytical models for calculating power and optimal sample size.

## Methods

### Data Simulation framework

MOPower simulates data based on statistical distributions. The framework is designed to consider a mixture of effects from each omics feature on the outcome of interest. Both single and multi-omics data can be generated. Figure 1 displays a flowchart of the current simulation framework, depicting the relationship between omics features and the outcome of interest.

**Figure. 1.**
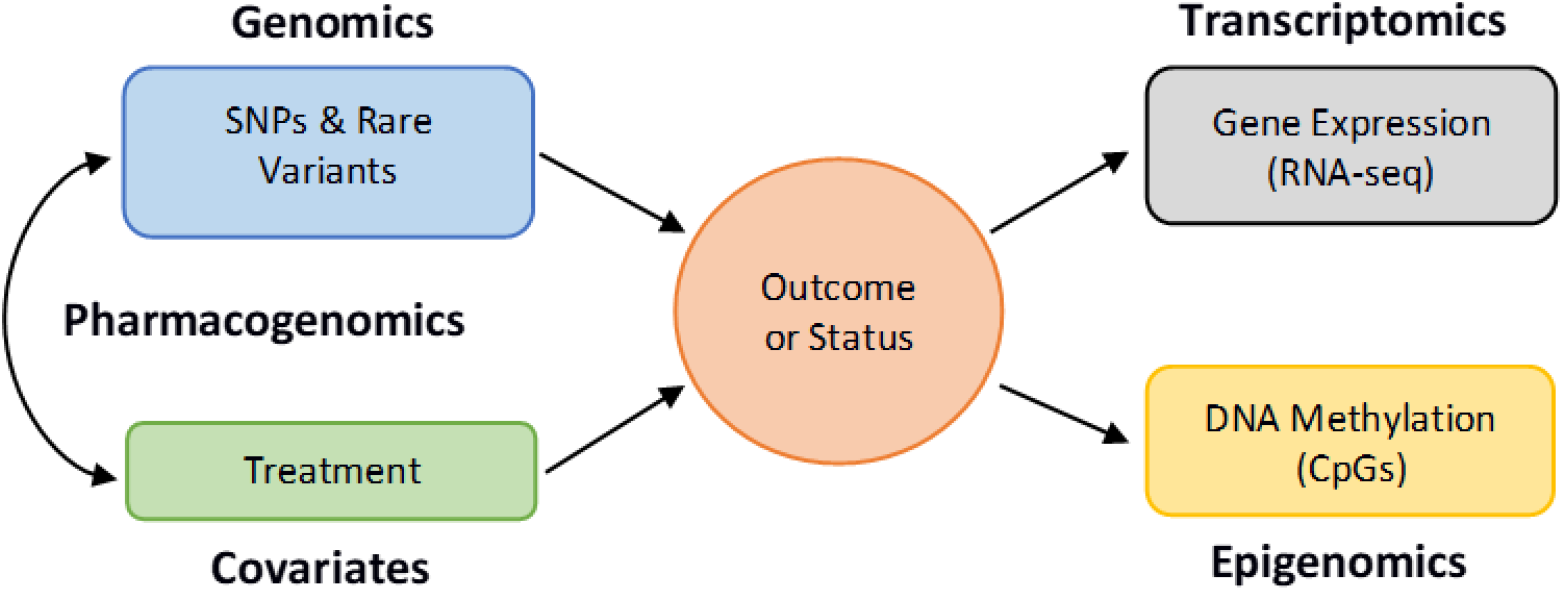
MOPower data simulation framework. Black lines represent the flow of information from simulation output to input processes.

### Outcome data

Users have the option of simulating a case-control setting or time-to-event outcome for each individual. Cases and controls are categorised for all individuals using user specification of the sample size and proportion of cases. A linear predictor model using a Binomial distribution assigns an individual a case or control status.

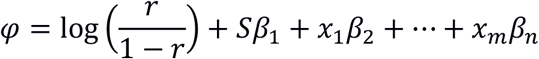

Where *φ*is the intercept, *r*is the case-control ratio, *S*is the set of single nucleotide polymorphisms (SNPs), and rare variants, the vector of *β*’s are effect sizes, and *x*represents additional covariates such as treatment. The response *Y*is, therefore:

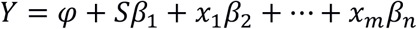

The probability of prevalence of the outcome status to be true is *π*=*e*^*Y*^/(1+*e*^*Y*^)for the Binomial random distribution. However, if genomic features are not included in the simulation, the case-control ratio is taken as the probability for the Binomial distribution.

For time-to-event data, a Weibull distribution is used to simulate survival times for each individual. MOPower implements only the most common type of censoring, which is ‘right censoring’. Right censoring occurs randomly during the study period and at the end of the study. To simulate survival times, the scale parameter of the Weibull function incorporates the SNPs and covariates into the simulation, similar to the case-control predictor model.

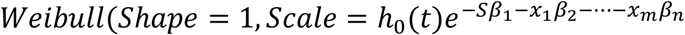

In this function, the baseline hazard, *h*_0_(*t*), is user-specified and controls the dispersion of the simulated times. The censoring indicator is classified as a percentage of the individuals for whom the event has occurred. This is introduced through a Binomial distribution. To determine the censored status of an individual, the survival time must be greater than the end of study time and the censoring indicator simulated with a binomial distribution must be 0. Every other combination is an event status for an individual. The observed outcome of an individual is created using the simulated event time (*E*), end of study time (*S*) and censoring indicator (*C*):

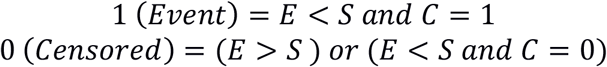

For a more detailed description of the simulation framework see Syed *et al*. [5].

### Multi-Omics data

MOPower simulates a select number of features from the genome, transcriptome and epigenome. SNP and rare variant data are simulated with a multinomial random distribution where the user specifies the range of possible minor allele frequencies (MAF). An additive model is assumed where 0 is the homozygous genotype, 1 is the heterozygous genotype, and 2 is the mutant homozygous genotype. The effect size of each SNP is determined by the MAF, and a user input baseline SNP effect range. This range is used to simulate a more realistic setting compared to a fixed effect size for all simulated SNPs. Rare variants are most often analyzed collectively through the creation of variant sets, burden or dispersion. MOPower will aggregate all rare variants in the set, creating a weighted burden.

Gene expression count data is simulated using the Negative Binomial (NB) distribution. The user is required to specify the *log*_2_fold change (effect size) between groups of individuals and the mean gene expression read count. The dispersion parameter is user input and controls the spread of the simulated values. The NB distribution is a natural choice over the Poisson distribution as it provides a better fit for RNA-Seq data by allowing an over-dispersion parameter to capture extra variability over the mean. The simulation of RNA-Seq data is adapted from Yu *et al*. [8]. This paper reviews all previous RNA-seq simulation and power calculation tools.

Methylation data are simulated as array hybridisation signals, as opposed to sequencing read counts, using a Beta distribution for each patient at each CpG site. The mean difference in methylation between each group should be specified. To add distortion to the analysis for a more realistic setting, hemi- and un-methylated sites can be introduced. A methylation odds ratio is calculated for the user to know the difference between groups of individuals. Where mean methylation is represented as *MM*, the odds ratio is:

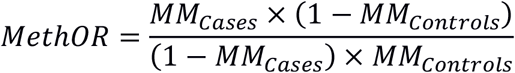

The simulation framework for DNA methylation is based on the work by Tsai *et al*. [12]. For most multi-omics feature data we implemented a random range of user specification values to mimic a realistic scenario because of the uncertainty in simulating more than one data type across a region in the genome, in contrast to simulating a specific effect size or dispersion for all features. Data simulation using a specific value of effect size or other input can still be achieved by setting the range to the same value. This is useful for single omics power calculation where you are only testing a single SNP, gene or CpG site over multiple simulations.

### Integration models

MOPower allows users to perform power calculation using some of the most recent and popular multi-omics analysis models. Some models find the correlation between omics features, and others combine the information into interconnected features, factors or clusters for further analysis. The majority of the methods require a regression model to be applied afterwards to obtain an analysis summary with p-values.

1. Joint regression modelling (Logistic and Cox proportional hazards regression [13]).
  - Regression modelling is commonly used to analyse multiple covariates with an outcome of interest. The logistic regression model caters for a binary outcome, and the Cox regression model is used for a time-to-event outcome. The user has two options for obtaining p-values from the regression model, either by i) aggregation of all features in each omics creating multiple sets or ii) fitting all features in the model and deducing significant features using a step-wise procedure.
2. Mediation analysis (Tingley et al. [14]).
  - Mediation analysis helps to explain the causal effects between omics features and an outcome of interest.
3. MOFA (Multi-omics Factor Analysis) (Argelaguet et al. [15]).
  - MOFA uses Bayesian and likelihood inference to disentangle the information from multi-omics data. The model correlates each feature with one another, creating a new matrix of factors. These factors are dropped one by one if their variance is lower than the users’ defined threshold.
4. Joint Non-negative Matrix Factorisation (NMF) (Taosheng et al. [16])
  - NMF is a type of integrative clustering analysis taken from the R package iClusterPlus. This method is very computationally intensive.
5. Similarity Network Fusion (SNF) (Wang et al. [17], Taosheng et al. [16]).
  - SNF is a network based analysis which captures both shared and complementary information from different data types. A network is created for each data type then combined into a similarity network. The data is integrated based on samples and not measurements.
6. GLM & Cox Path with L1 penalty (Park et al. [18], Mankoo et al. [19]).
  - This is a path following algorithm with an L1 regularisation procedure for variable selection. This algorithm implements the predictor-corrector method to determine the entire path of the coefficient estimates as the amount of regularisation varies.
7. Exact Test (Robinson et al. [20] and McCarthy et al. [21])
  - The exact test computes gene-wise tests for differences in the means between two groups of negative-binomially distributed counts.
8. NB Regression Model (Robinson et al. [20] and McCarthy et al. [21])
  - The NB generalized log-linear model is fit to the read counts for each gene. Additional covariates can also be fit with this model.

### Power calculation procedure

Each data replicate is fitted with an integration model to obtain test statistics. This is interpreted as the discovered association incorporating a mixture of genetic variant, gene expression and methylation data to identify the significance of a locus with the outcome of interest. Inferences can also be made on the relationship or interaction between different omics features. The statistic of importance from each analysis is the *p*-value. Power is then equal to the percentage of replicates for which *p* < type I error rate, for the total number of simulations.

#### Procedure

1. The user specifies all input parameters; this includes the number of simulations, initial sample size, number of genes or variants, mean gene expression, fold change between groups and the type I error rate. All input parameters are described in Table 1.
2. Data is simulated from statistical distributions using the input parameters.
3. Data is fit with integration and statistical models to obtain the test statistics under both the null and alternative hypotheses.
4. Power is calculated for the specified input parameters. Power is equal to the percentage of replicates *ρ*for which *p*< type I error rate.

**Table 1.**
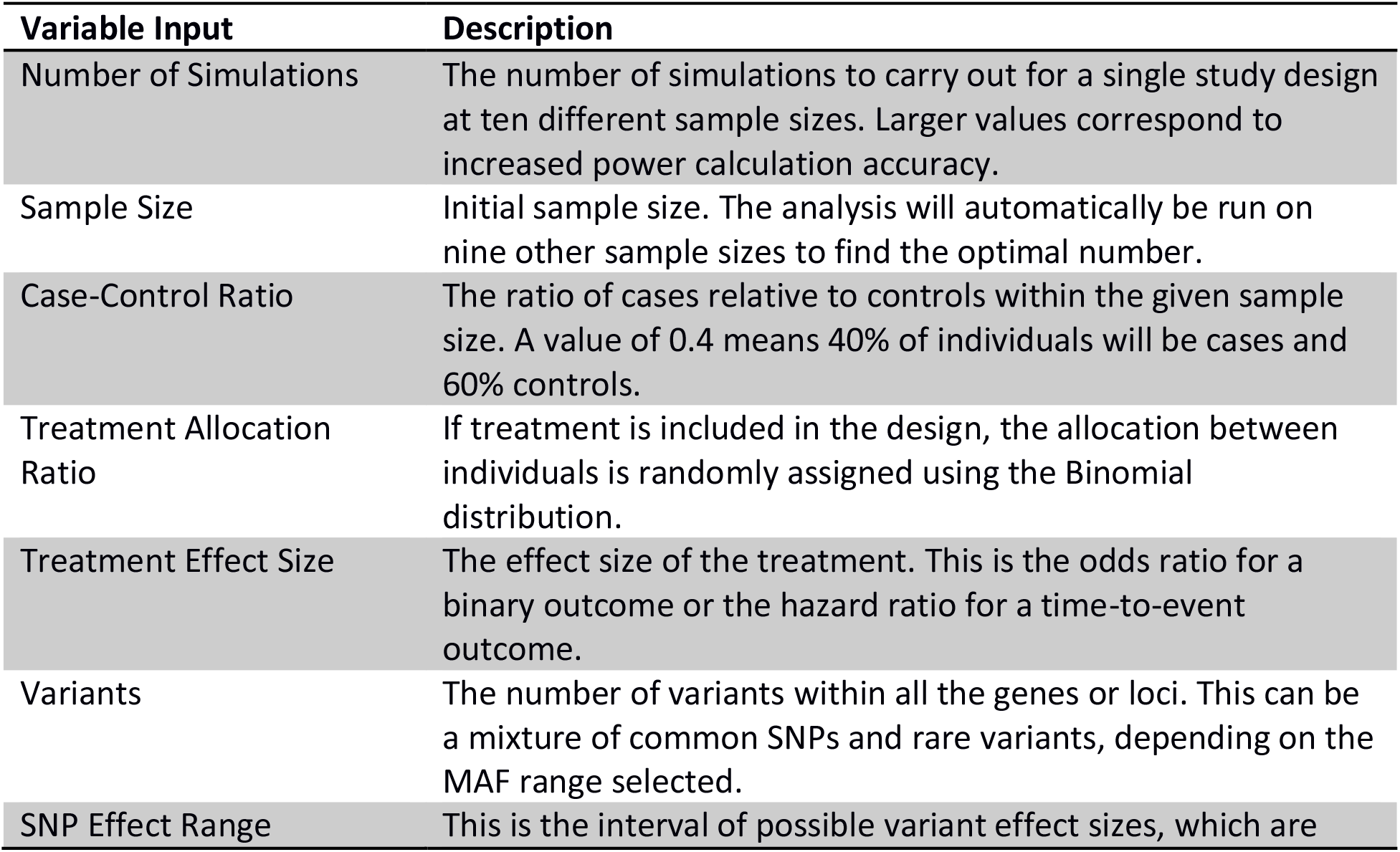
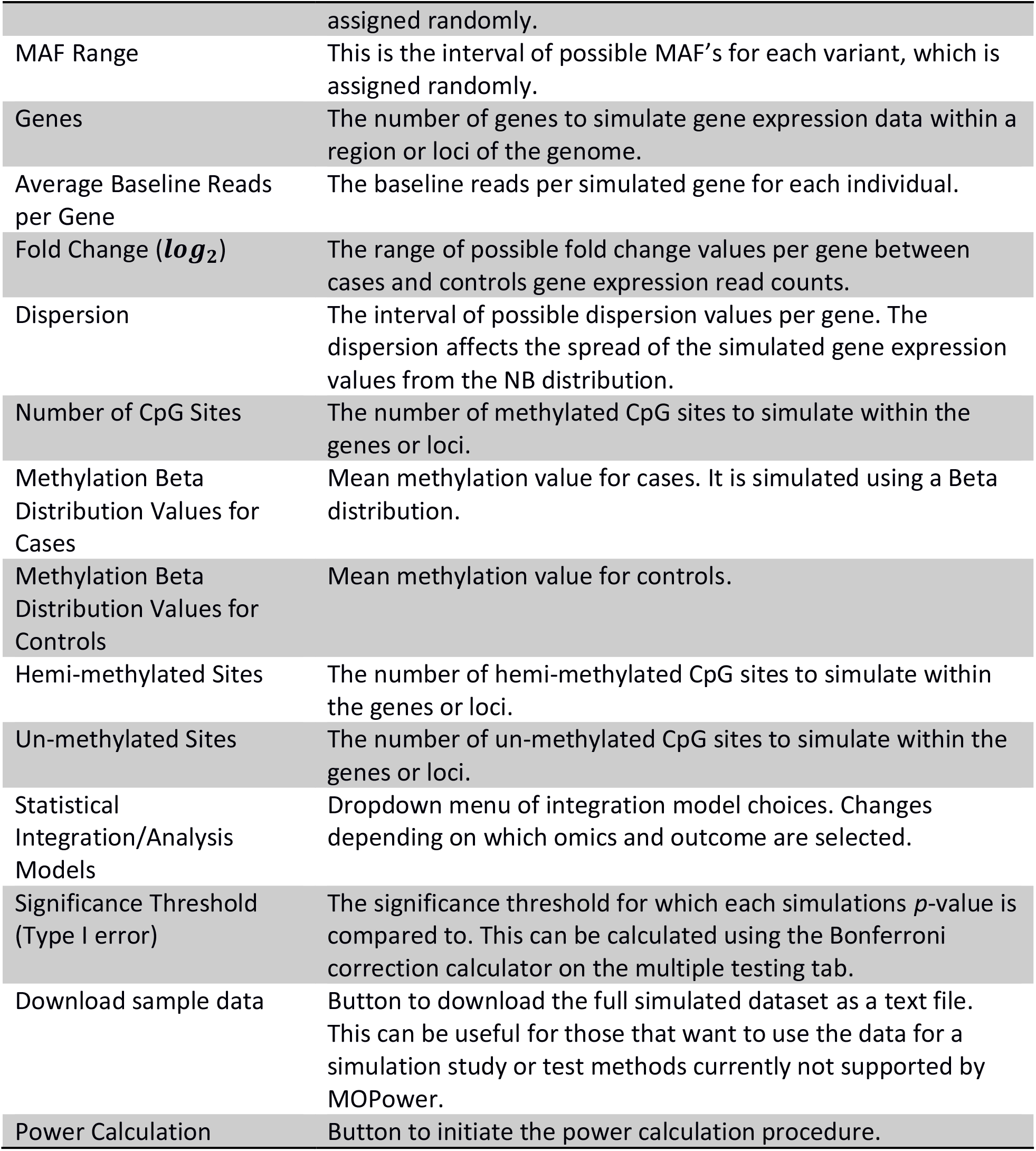
MOPower inputs, features and buttons description.

This is the same process for both single and multi-omics options with exemption to RNA-seq power calculation using the exact test or NB regression model from the *edgeR* package because these analyses reflect the likelihood of detecting differentially expressed genes within a given sample of genes, not over multiple simulations.

The power is displayed both in numerical and graphical form based on ten different sample sizes, including the user-specified sample size. On a separate tab, the false discovery rate (FDR) is calculated using the Benjamini-Hochberg procedure [22]. This informs us of the threshold for controlling the type I error rate in the presence of multiple testing. It is a less conservative procedure compared with the Bonferroni correction, which is also displayed on the same tab.

## Implementation

MOPower is a user-friendly R-shiny web application that is entirely interactive. No coding experience is needed, just an understanding of multi-omics experiments. MOPower can be launched through R studio and from any web browser, with the app hosted on the Shiny global server. If run locally, all package dependencies have to be downloaded and installed before running the application. A script is provided to facilitate this process. MOPower utilizes multi-processing using the packages *doparallel* and *foreach* to optimize efficiency when performing thousands of data simulation and analysis tasks concurrently. Figure 2 displays a process flowchart of the power calculation procedure from inputs to options for downloading data and displaying power plots. This figure shows how simple MOPower is to use, requiring very little interaction from the user.

**Figure. 2.**
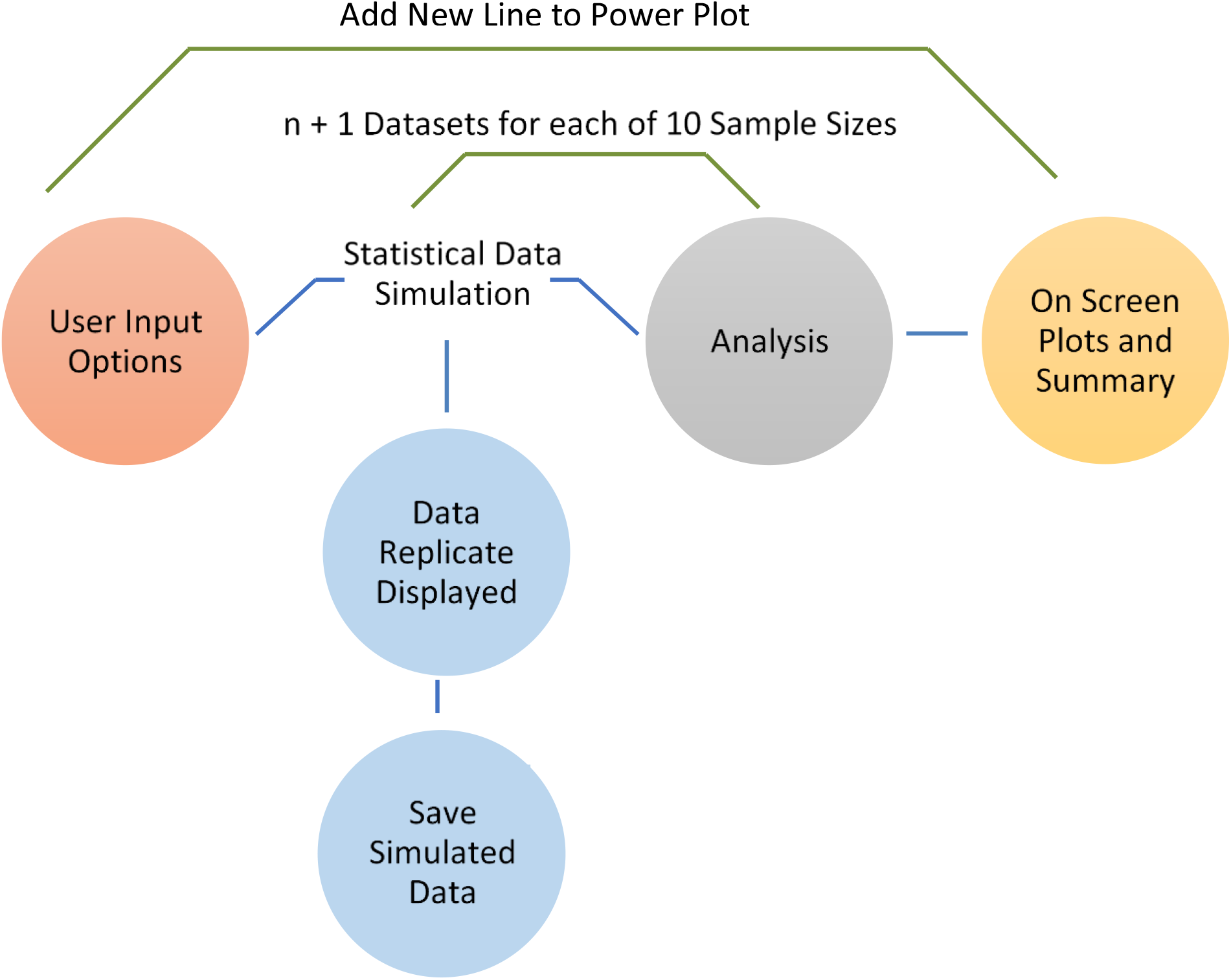
MOPower navigation and process flowchart. Blue lines represent the flow of information from left to right. Green lines represent the flow of information from right to left.

### User interface

The MOPower user interface is made up of seven tabs; i) the power calculator; ii) multiple testing; iii) instructions; iv) about; v) news and updates; vi) frequently asked questions and vii) references. When using MOPower useful tooltips and messages appear throughout to guide the user, as shown in Figure 3. In this example, the user is provided with information on performing RNA-seq power calculation using the exact test or NB model. It is initiated and displayed when the user selects one of these methods and checks the RNA-seq checkbox.

**Figure. 3.**
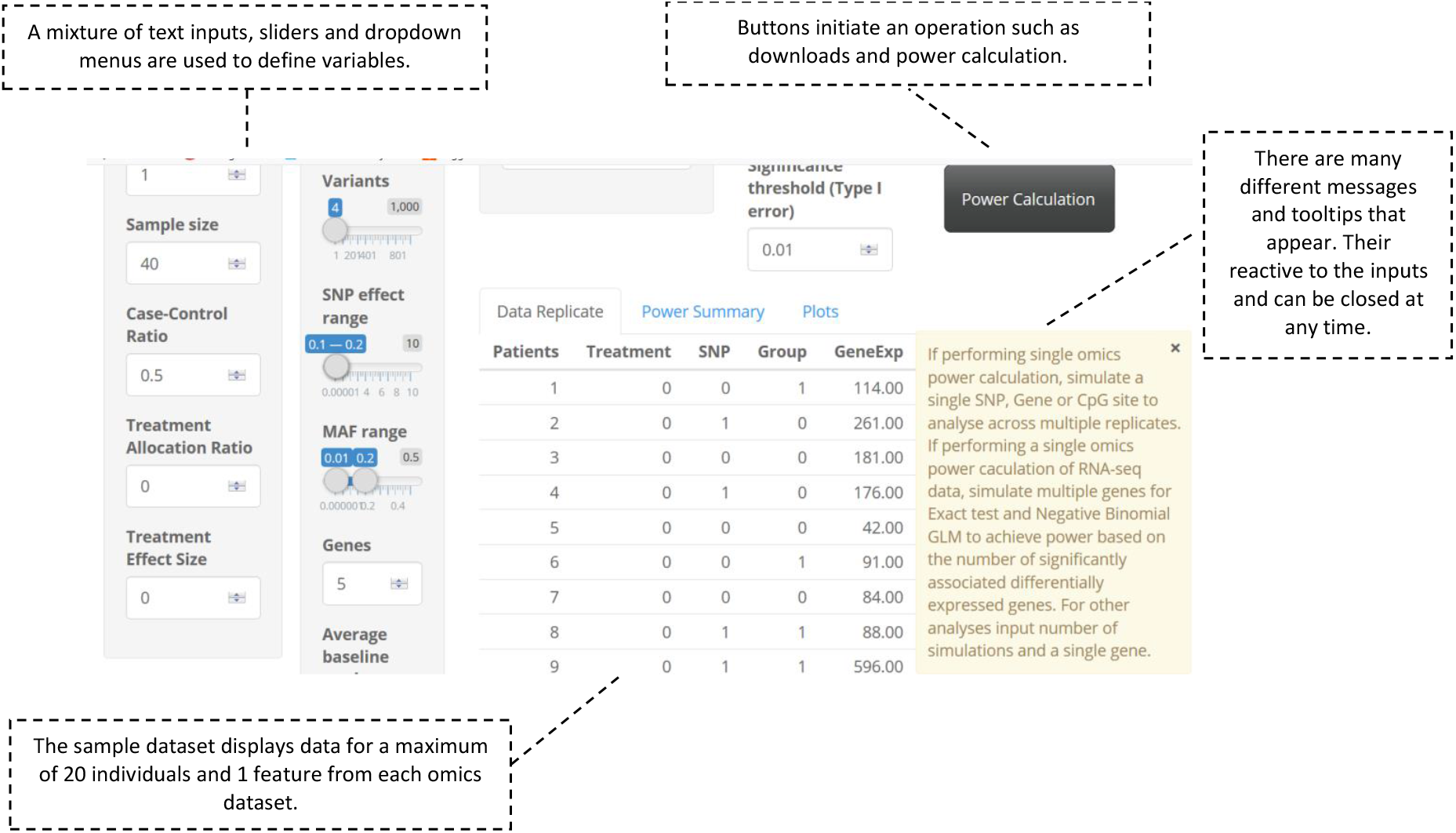
MOPower interface showing warning message example, reactive data sample and inputs. Warning messages and tooltips are automatically displayed with specific user inputs and selections.

Input options and selections are essential to simulating realistic data. MOPower offers several omics features for users to customize their study design based on genomic, transcriptomic or epigenomic data. Table 1 describes all inputs, selections and buttons. Currently, only a single feature per omics is implemented, but with multiple adjustable factors such as effect sizes and the number of genes, variants and CpG sites.

MOPower produces a variety of different outputs on-screen. First, a sample dataset for the first 20 individuals and one variable from each omics feature is shown which interactively changes as the user adjusts input parameters. The sample data can be used to access if the correct data has been simulated before power calculations can begin. Second, after the power calculation has finished an FDR plot to assess the inflation of the type I error rate and a histogram of distributed p-values are displayed on the multiple testing page. Third, the power for each sample size run is displayed numerically in a table and as a graph. Figure 4 depicts a time-to-event setting power calculation using the CoxPath model. The plot can be downloaded in Portable Network Graphic (.png) format and can be manipulated using zooming and panning controls. Also, there is an option to add the next power calculated line to the same plot, which is helpful for direct comparison of methods or different study design parameters.

**Figure. 4.**
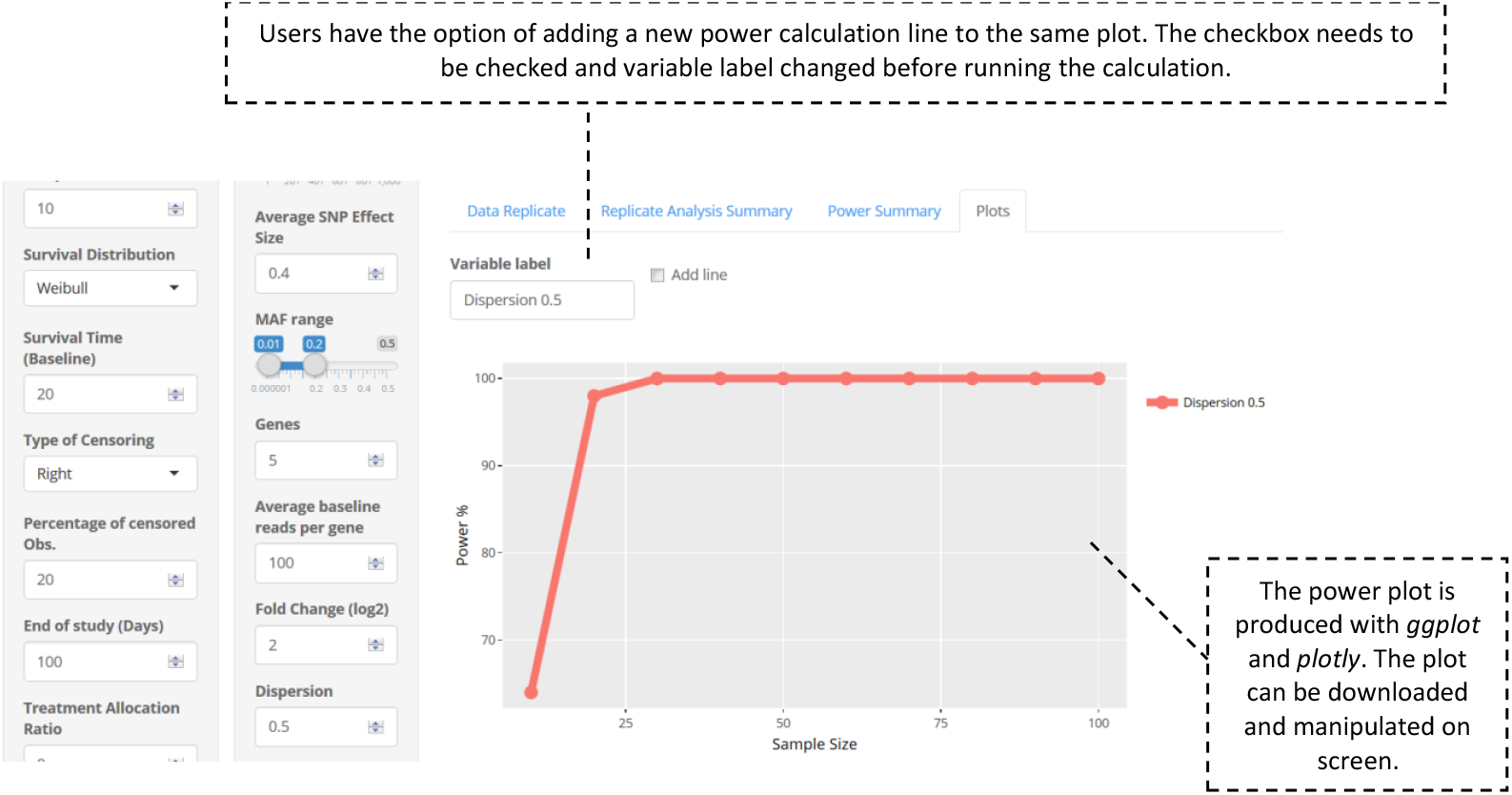
MOPower interface. This example depicts output from a time-to-event analysis using the CoxPath method, testing the power to detect an association with SNP, gene expression and methylation data for an initial sample size of 20 individuals.

## Results

### Simulation study

MOPower has been evaluated through a simulation study testing a single scenario with multiple analysis models. The following results are a presentation of the use of the software and not a detailed comparison of multi-omics scenarios and analysis models. This study not only looked at the performance of the software but also estimated optimal sample size for the scenario and evaluated the power to detect an association with a binary outcome with varying number of omics features for a given loci using several integration models. The study design we evaluated is a case-control design where 15 individuals are split evenly between case-control groups with SNP genotype and gene expression count data on each individual. The locus of interest contains 20 SNPs with a varying MAF range of common variation between 0.05 and 0.5. The SNP effects also vary randomly from 0.1 and 1, which is a moderate effects range. Gene expression data have been collected on five genes with an average baseline reads of 100 with the fold change between cases and controls in the range 1 to 2. The dispersion values are between 0 and 2. 100 simulations were run, allowing for a Bonferroni corrected threshold for each analysis based on the number of statistical tests performed (0.0001). Figure 5A illustrates significant discrepancies in power between the integrative models. The logistic regression model has the largest power to detect an association with the optimal sample size of 105 individuals. Mediation analysis achieves at least 80% power with a sample size of 140, with SNF achieving a peak of 73% power with a sample size of 135. MOFA has a sharp decrease in power to 0% after a sample size of 30. For this scenario, a sample size between 100 and 150 would be optimal, except for MOFA and NMF which have inflated type I error rates and a distinct lack of power to detect an association at that sample size. We next asked how including a larger number of omics features influences the power calculations (Figure. 5B and 5C). As a result, some methods (Joint Regression and MOFA) increase in power, while others (SNF and mediation analysis) decrease in power when the number of omics features increases. MOFA achieves greatest power when the sample size number corresponds to the number of features included. More features can distort results of some analyses or can validate the evidence of association in other analyses. A concerning result was the lack of any power for NMF even with increasing sample size. This is likely the result of a stringent type I error rate used for each analysis. The volatility of power for each analysis is due to the randomness of the data at each replicate. The range of values is different for every run and can cause inaccuracy. To avoid this, smaller intervals for values such as MAF and effect sizes would increase the probability of a steady increase in power with sample size.

**Figure 5.**
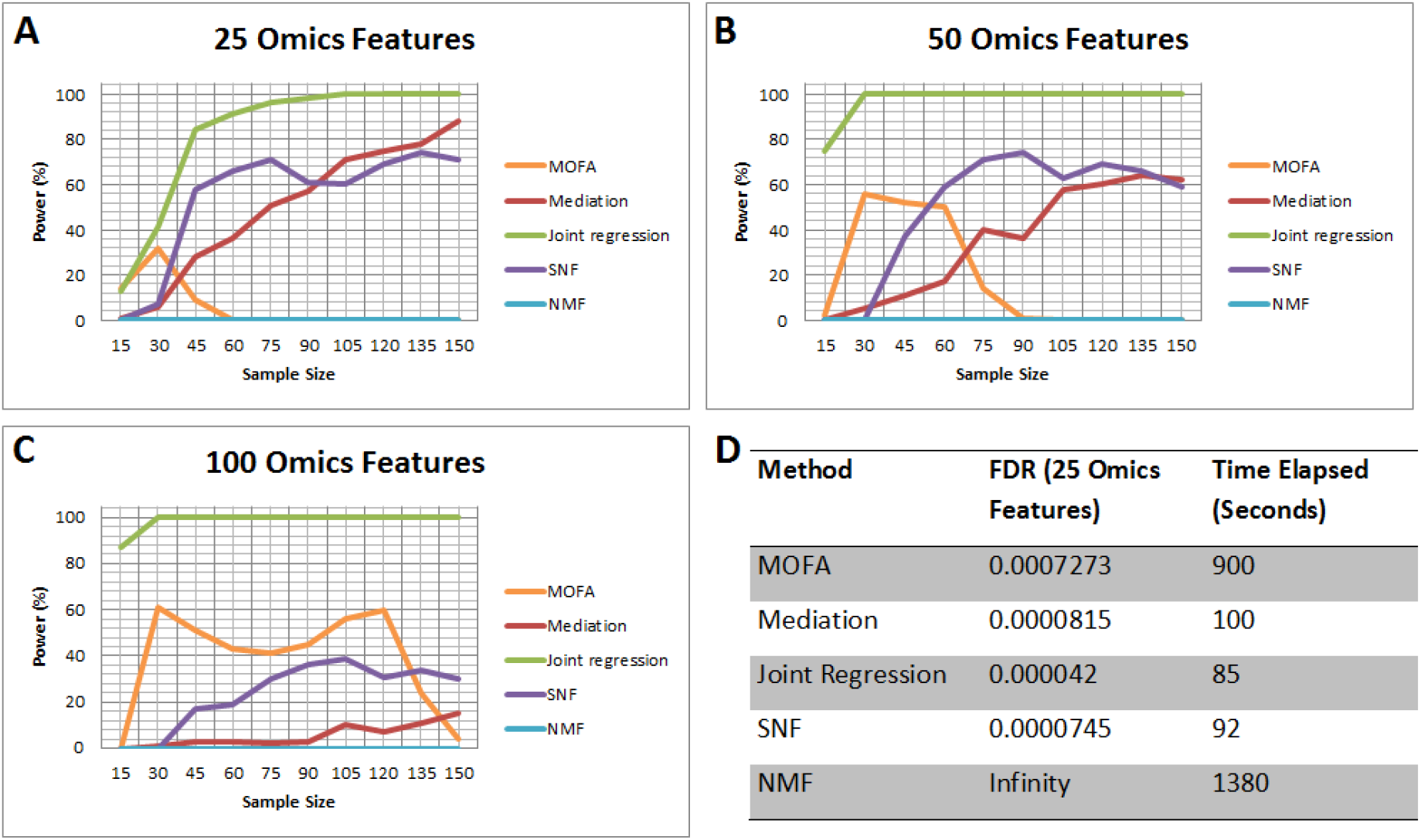
Simulation study power calculation plots and table. A: Power plot of 20 SNPs and expression of 5 genes for five integrative analyses with a sample size ranging from 15 to 150. B: Power plot of 40 SNPs and expression of 10 genes for five integrative analyses with a sample size ranging from 15 to 150. C: Power plot of 25 omics features for five integrative analyses with a sample size ranging from 15 to 150. D: Summary table of false discovery rate and time elapsed for analysis using the five integrative models.

Figure 5D indicates that the FDRs for all models are small and that MOFA and NMF are the most computationally demanding methods to use. Time for each power calculation to finish can differ greatly dependent on integration model, but the most influential factor affecting processing speed is the number of available cores on a computer or cluster that the software can access. The use of one model over another should be carefully considered in terms of the interpretation benefits of the results produced and under which circumstances the model performs best.

## Conclusions

Multi-omics studies have the potential to change our understanding of the biological mechanisms that underlie many biological and physiological processes. MOPower is the first multi-omics data simulator and power calculator and will aid investigators undertaking multi-omics experiments by providing information on study design, appropriateness of an integration model, power and sample size to achieve the optimal results from their study. This paper has demonstrated the use of MOPower through a simulation study which identified the optimal sample size and integration model for one particular multi-omics setting. MOPower has the framework available to become more sophisticated by offering other omics study design parameters from proteomic and metabolomics data. Furthermore, new integration methods can be implemented to give investigators a broader choice, such as generalised linear mixed-effects regression modelling, Bayesian mixed models [23], sparse partial least squares regression [24], matrix integration [25, 26] and machine learning models [27] where we can measure power in terms of prediction accuracy. Deep-learning models have already been successful in predicting survival in liver cancer patients using multi-omics data [28]. We expect the availability of a framework for power calculation like MOPower to be of great benefit both in the initial design, and during the implementation phase of such studies.

## Availability and requirements

**Project name:** MOPower

**Home page:** MOPower is available for download at https://github.com/HSyed91/MOPower and is available to launch from https://hsyed.shinyapps.io/MOPower/

**Operating system(s):** Linux, Windows, MAC OSX

**Programming language:** R, Python

**License:** GNU (General Public License) v3

**Any restrictions to use by non-academics:** None

## Abbreviations

SNPs: Single nucleotide polymorphisms
MAF: Minor allele frequency
NB: Negative Binomial
NMF: Non-negative Matrix Factorization
MOFA: Multi-omics Factor Analysis
SNF: Similarity Network Fusion
FDR: False discovery rate.

## Acknowledgements

The authors wish to thank the entire GOSgene team at the Institute of Child Health UCL, for their assistance in testing and providing feedback on MOPower.

## Authors’ Contributions

HS carried out the literature review on power calculation and data simulation for SNP, RNA-seq and methylation data. Commonly used software for individual omics was also covered in the review. HS also coded, developed, debugged and tested the software and drafted the manuscript. PB and DK conceived the general project and supervised it. GWO, DK, CB and PB participated in the design and coordination of the software and helped to draft the manuscript. All authors read and approved the final manuscript.

## Ethics approval and consent to participate

Not applicable.

## Consent for publication

Not applicable.

